# Sorbitol in loquat promotes early flower bud differentiation via the MADS-box transcription factor EjCAL

**DOI:** 10.1101/2021.08.17.456699

**Authors:** Hongxia Xu, Ting Chen, Meng Qi, Xiaoying Li, Junwei Chen

## Abstract

The sugar alcohol sorbitol plays an important signaling role in fruit trees. Here, we found that sorbitol significantly increased during flower bud differentiation (FBD) in loquat (*Eriobotrya japonica* Lindl.) from the physiological FBD stage (EjS1) to the morphological FBD stage (EjS2), and it then decreased in the panicle development stage (EjS3) compared to in EjS2, and in subsequent stages. Spraying sorbitol increased the sorbitol content and thereby promoted early FBD and increased the proportion of flower buds that completed FBD. A transcriptomics analysis showed that the expression of a MADS-box transcription factor (TF) family gene, *EjCAL*, was highly correlated with the FBD phenotypic data. *EjCAL*-overexpressing transgenic tobacco exhibited the early FBD phenotype. Using the *EjCAL* promoter as bait in a yeast-one hybrid (Y1H) assay, the TF ERF12 was identified. Chromatin immunoprecipitation (ChIP)-PCR confirmed that EjERF12 can bind to the *EjCAL* promoter, and β-glucuronidase (GUS) activity assays demonstrated that EjERF12 can regulate *EjCAL* expression. Spraying loquat with sorbitol confirmed that *EjERF12* and *EjCAL* expression were regulated by sorbitol. We also identified downstream functional genes (*EjUF3GaT1, EjGEF2*, and *EjADF1*) that might be involved in FBD. Finally, we found that the change in the level of hyperoside (a reproduction-related flavonoid) was consistent with that of sorbitol during FBD in loquat, and EjCAL can bind to the *EjUF3GaT1* promoter and might thereby regulate hyperoside biosynthesis. Two early- and late-flowering varieties of loquat and *EjCAL*-overexpressing transgenic tobacco plants were used to confirm this hypothesis.

**One-sentence summary:** Sorbitol promotes bud differentiation via EjCAL.

## Introduction

Flower bud differentiation (FBD) is a complex morphogenesis process that has a direct impact on the quantity and quality of flowers as well as the fruit set rate, which affects the yield. FBD can be divided into the physiological FBD (EjS1) and morphological FBD (EjS2) stages. It is influenced by interactions among environmental factors (such as photoperiod and temperature) (Song et al., 2013) and internal factors (such as carbohydrates and hormones) (Bernier et al., 1993; Hanke et al., 2007; Turnbull, 2011). Sugar is an important carbon source and energy substance in plants and also regulates flower bud growth and development (Ito et al., 2002, 2004; Xing et al., 2015). Recent research has shown that trehalose-6-phosphate (T6P) is an indicator of carbohydrate in plants, and *T6P synthase* (*TPS*) is a key gene in its biosynthesis pathway (Vandesteene et al., 2012) Additionally, in *Arabidopsis thaliana*, in response to sugar signaling, AtTPS1 induces flower formation by upregulating key flowering genes such as *LEAFY* (*LFY*) and *APETALA1* (*AP1*) (Wahl et al., 2013). Soluble sugars such as sucrose and glucose have been shown to be critical in the regulation of flower formation. Applying sucrose to *Vitis vinifera* upregulates *TERMINAL FLOWER 1* (*VvTFL1*)and promotes the formation of apical flower buds (Yang et al., 2011). In apples, sucrose plays an important signaling role in flower bud induction by regulating the expression of *SUPRESSOR OF CONSTANS 1* (*SOC1*), *AGAMOUS like 24* (*AGL24*), and other flowering genes (Xing et al., 2015). In tobacco, floral organs can still develop normally when exogenous glucose is applied under dark conditions, but if the glucose concentration is reduced, no floral organs are produced, or abnormal development occurs (Cousson and Van, 1983). However, glucose has a dual effect in the regulation of flower formation, as after culturing *A. thaliana* seedlings with 2% or 6% glucose, the latter resulted in delayed flowering of mosses compared to the former, regardless of the sunshine exposure duration (Rolland et al., 2006).

Sorbitol is a special photosynthetic product that is produced by many economically important fruit trees in the Rosaceae family. It accounts for 60–80% of the photosynthates produced in leaves and transported via phloem (Bieleski et al., 1982; Loescher, 1987; Cheng et al., 2005). Sorbitol can also be transformed into fructose and glucose through physiological pathways, and the conversion rate of sorbitol is higher than that of sucrose (Morandi et al., 2008). Sorbitol plays various roles in stress resistance and growth and development (Zhou et al., 2003; Kanayama, 2009). There are increasing numbers of studies on the role of sorbitol in flower development, and sorbitol is thought to be a signaling factor that regulates this process (Archbold, 1999). However, the role of sorbitol in FBD regulation in fruit trees is unclear and rarely studied.

Floral initiation is under tight genetic control (Willmer and Riechmann 2010). *LFY, AP1*, and the *AP1* paralog *CAULIFLOWER* (*CAL*) have been identified as factors that control the onset of flower development (Goslin et al., 2017). In *Arabidopsis, AP1* and *CAL* have been found to be duplicate genes generated via a whole-genome duplication event in the flowering plant family Brassicaceae (Wang et al., 2012). Silencing of *CAL* alone does not cause any obvious phenotypic change, while silencing of *CAL* in the *ap1* mutant background enhanced the phenotype of the plant, suggesting that *CAL* and *AP1* may have redundant functions (Bowman et al., 1993). Using genetically modified *Arabidopsis* overexpressing *LFY* or *AP1*, it was shown that these genes are sufficient to promote flower initiation and development (Weigel and Nilsson, 1995; Mandel and Yanofsky, 1995). After constitutive overexpression of *AP1* from *Arabidopsis*, early flowering was obtained in transgenic citrus (Peña et al., 2001) and tomato (Ellul et al., 2004). Additionally, *FRUITFULL* (FUL), which is expressed in the shoot apical meristem at the transition to flowering, may also act as a floral integrator (Schmid et al., 2003). Similarly, *MADS-box transcription factor 5* (*MdMADS5*) from apple (*Malus domestica*) transformed into *Arabidopsis* also promoted FBD, resulting in early FBD (Kotoda et al., 2002). However, other MADS-box genes such as *FLOWERING LOCUS C* (*FLC*), *TFL1*, and *SHORT VEGETATIVE PHASE* (*SVP*) have negative regulatory roles in flower development (Boss et al., 2004). Constitutive *MdTFL1* overexpression under the control of the cauliflower mosaic virus promoter (*CaMV35S*) in *Arabidopsis* slows the transition from the vegetative to reproductive phase (Kotoda and Wada, 2005). Further, *MdTFL1* downregulation in transgenic apple shortens the juvenile stage and induces early flowering (Kotoda et al., 2006).

Loquat (*Eriobotrya japonica* Lindl.) belongs to the Maloideae subfamily of the Rosaceae family. It is a tropical and subtropical evergreen fruit tree (Lin et al., 2010). In southeast China, physiological FBD in loquat begins in midsummer, and its flowers bloom from October to February. The fruits originating from the early flowers that bloom from October to November are usually large and of good quality, but the fruits originating from the later flowers are vulnerable to low temperature and freezing damage in the northern subtropical region (Xu et al., 2017). Therefore, FBD and the blooming time affect the fruit yield and quality. Although several genes related to flower development in cultivated loquat, such as *EjLFY, EjAP1, EjFT, EjSOC, EjTFL1*, and *EjAGL17* have been cloned (Esumi et al., 2005; Liu et al., 2013; Reig et al., 2017; Jiang et al., 2019; Jiang et al., 2020; Xia et al., 2020), the mechanism of FBD in loquat remains unclear.

Here, we aimed to identify the metabolic factors and gene networks controlling the initiation of loquat FBD. We demonstrated that sorbitol signaling plays an important role in inducing FBD in loquat, and a MADS-box transcription factor (TF) designated *EjCAL* was significantly related to FBD. Chromatin immunoprecipitation (ChIP)-PCR and yeast one-hybrid (Y1H) assays confirmed that EjERF12 can bind to the promoter of *EjCAL* and regulate its expression, and both genes were regulated by sorbitol. We also found that sorbitol promoted the accumulation of an important developmental flavonoid, hyperoside, and demonstrated that EjCAL can bind to the *EjUF3GaT1* promoter and might thereby regulate hyperoside biosynthesis. This study provides a new viewpoint and evidence on the regulation of FBD by sorbitol signaling in loquat.

## Results

### Sorbitol was significantly accumulated during FBD in loquat

Flower development in loquat can be divided into five stages based on the morphological and histological changes, designated EjS1 to EjS5 (Figures 1A–J). EjS1 was not significant period of FBD, which designated physiological FBD (Figures 1A and 1F), EjS2 was the significant period of FBD, which designated the morphological FBD stage (Figures 1B and 1G) and EjS3 was the panicle development stage (Figures 1C and 1H). In EjS2, the apical meristem (AM) of EjS1 transformed to the flower meristem (FM). In EjS3, it gradually formed a floral structure, including a petal primordium (PEP), stamen primordium (STP), and pistil primordium (PIP).

**Figure 1.**
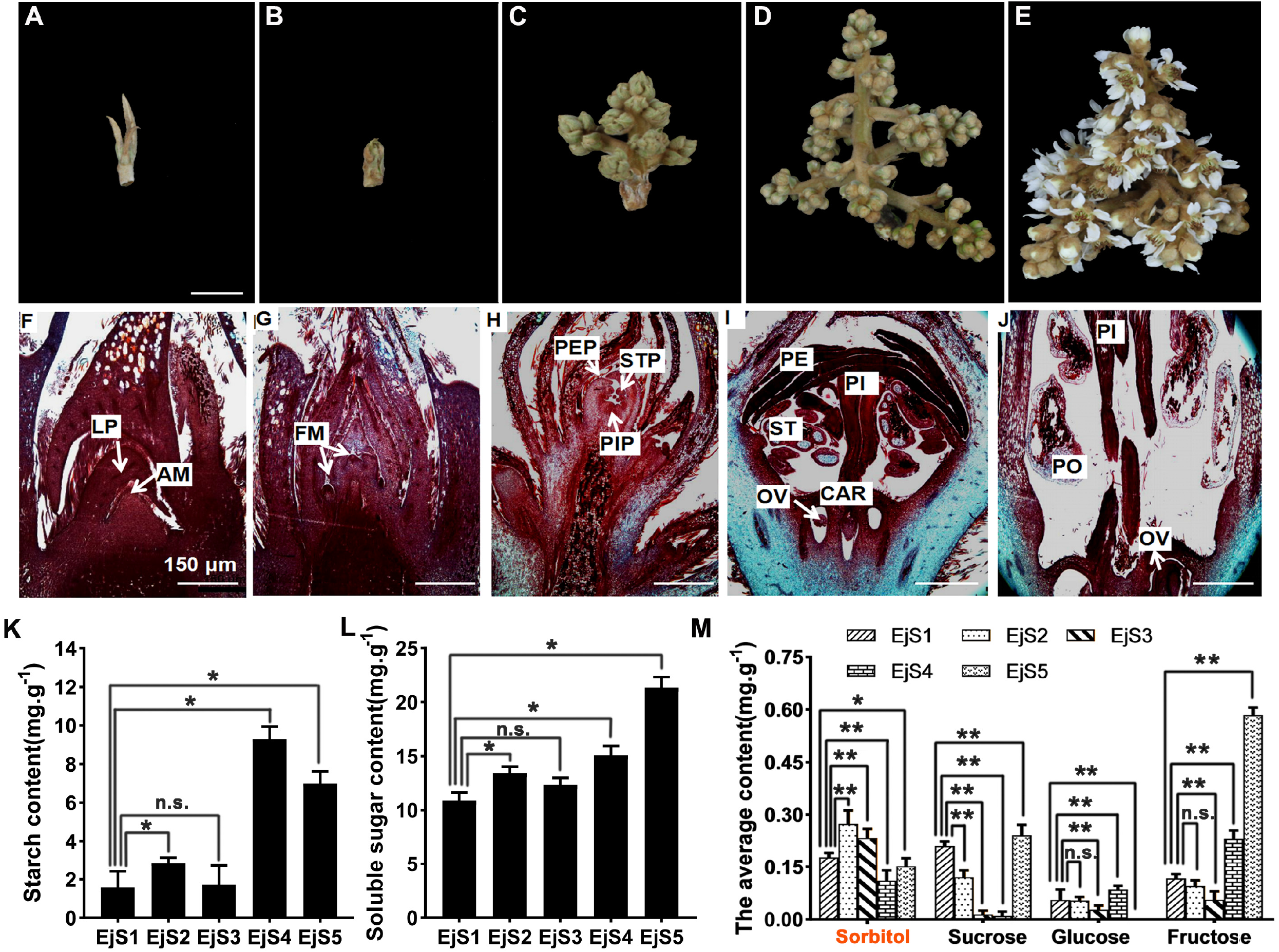
Flower bud differentiation (FBD) and changes in carbohydrate content in loquat. **A–E,** Five stages (EjS1 to EjS5) of the flower development process in loquat. **F,** EjS1 stage: leaf primordium (LP) and apical meristem (AM). **G,** EjS2 stage: flower meristem (FM) structure. **H,** EjS3 stage: petal primordium (PEP), stamen primordium (STP), and pistil primordium (PIP). **I,** EjS4 stage: petal (PE), pistil (PI), stamen (ST), and ovary (OV). **J,** EjS5 stage: pistil (PI), pollen (Po), and ovary (OV). **K–L,** Starch and soluble sugar content from EjS1 to EjS5. **M,** Sucrose, glucose, fructose, and sorbitol content from EjS1 to EjS5. Data are expressed as mean±SEM of n=3 biological replicates. *P<0.05, **P<0.01 (Student’s t-test).

Based on the above morphological data, the starch and soluble sugar content in the five stages were measured (Figures 1K and 1L). There were obvious increases in both starch and soluble sugar content from EjS1 to EjS2 during FBD. Therefore, we further examined the content of four important sugars (sucrose, glucose, fructose, and sorbitol) in the five stages (Figure 1M). The glucose and fructose content did not change from EjS1 to EjS2. The sucrose content continuously decreased from EjS1 to EjS3. Critically, the sorbitol content increased in EjS2 and then decreased in EjS3, which was consistent with the FBD phenotypic data, and the sorbitol content was much higher than the other three sugars in EjS2 and EjS3. The results indicate that sorbitol accumulation might be related to the AM transformation into the FM.

### Spraying sorbitol promoted loquat FBD

To assess the function of sorbitol in FBD, 1.0% sorbitol was sprayed on the loquat plants prior to FBD in July. After 10 days, the control buds were still in the EjS1 stage, while the sorbitol-treated buds had developed into the EjS2 stage (Figure 2A). After 40 days, The sorbitol content in the flower buds was significantly upregulated after both 10 and 40 d of treatment compared to in water-treated control buds (Figure 2B). After statistical development period, only about 20% of the control buds developed to the EjS2 stage, while 25% of the sorbitol-treated buds developed to the EjS2 stage and 15% developed to the EjS3 stage (Figure 2C). After 40 days of sorbitol treatment, the ratio of EjS2–EjS5 stage buds/total buds was about 40%, which was higher than after control treatment (20%) or other sugar treatments (Figure 2D). These results indicate that sorbitol accelerated FBD. We also assessed the ratio after 80 days of treatment. The ratio was still higher after sorbitol treatment than after control or other sugar treatments, but the gap was closing (Figure 2E). This indicated that sorbitol can accelerate FBD and significantly increase the proportion of flower buds in FBD.

**Figure 2.**
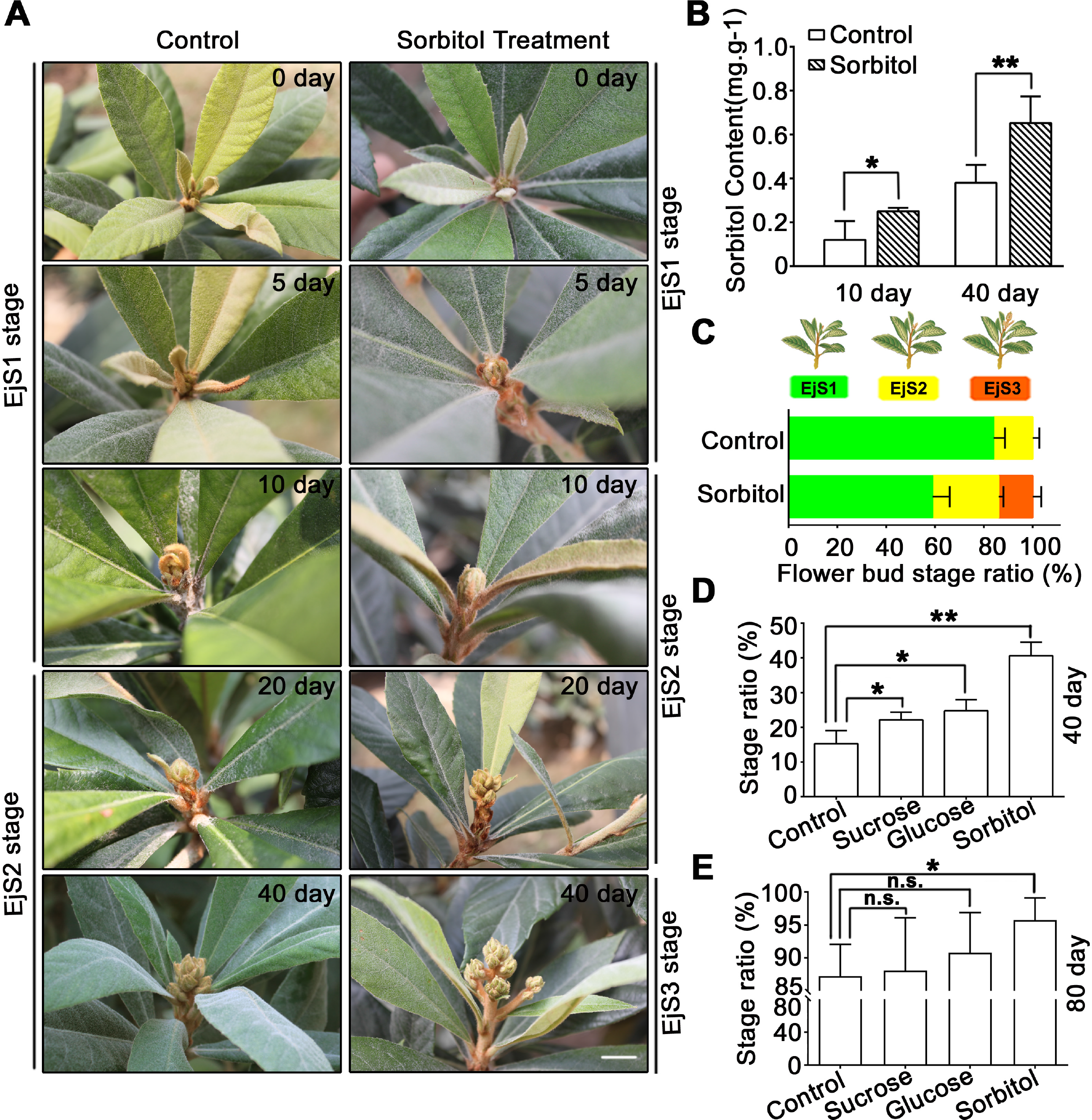
Flower bud differentiation (FBD) after spraying sorbitol. **A,** The EjS2 stages occurred after spraying water for 20 days, while the EjS2, and EjS3 stages occurred after spraying sorbitol for 10, 20, and 40 days. **B,** Sorbitol content in control and sorbitol groups after spraying for 10 and 40 days. **C,** Flower bud stage ratios (EjS1, EjS2, and EjS3 stage buds/total buds) in control and sorbitol groups after spraying for 40 days. **D,** Flower bud stage ratios (EjS2–EjS5 stage buds/total buds) when spraying sucrose, glucose, or sorbitol after 40 days. **E,** Flower bud stage ratios (EjS2-EjS5 stage buds/total buds) when spraying sucrose, glucose, and sorbitol after 80 days. Data are expressed as mean±SEM of n=3 biological replicates. *P<0.05, **P<0.01 (Student’s t-test).

### TF EjCAL was related to FBD based on RNA-seq data

To identify the molecular mechanism underlying FBD, flower buds in the five FBD stages were subjected to RNA-seq. We focused on the early stage of FBD, when the sorbitol content first increased (EjS2 stage) and then decreased (EjS3 stage) (Figure 1). The expression of candidate genes (based on RNA-seq data) had to meet all of the following criteria: EjS2 > EjS1 (474 genes), EjS2 > EjS3 (559 genes), EjS3 > EjS1 (3753 genes), EjS3 > EjS4 (8385 genes), and EjS3 > EjS5 (11195 genes) (Figures 3A and 3B) (i.e., the relative expression is higher in EjS2 than EjS1/EjS3/EjS4/EjS5, and in EjS3 than EjS1/EjS4/EjS5). We identified five genes that fulfilled these criteria; one TF (designated EjCAL) and four functional genes (including genes that encode laccase-14 like protein and cysteine-rich repeat secretory protein 55). TFs serve as upstream switches and regulators, so we focused on the function of the single TF (Figure 3C). Its expression was highest in the EjS2 stage (five times higher than in the EjS1 stage) (Figure 3D). We analyzed its sequence and found that it had a typical MADS-box domain (Figure 3E). The neighbor-joining tree analysis indicated its similarity to AtMADS79 and showed that it belongs to the MADS-box TF family. It has one 60-aa MADS-box domain in the N-terminus and a coiled-coil domain and a low-complexity domain in the C-terminus (Figure 4B). To confirm its basic role as a TF, we analyzed its subcellular localization in tobacco leaves. The results showed that it was expressed in the nucleus and can function as a TF (Figure 4C).

**Figure 3.**
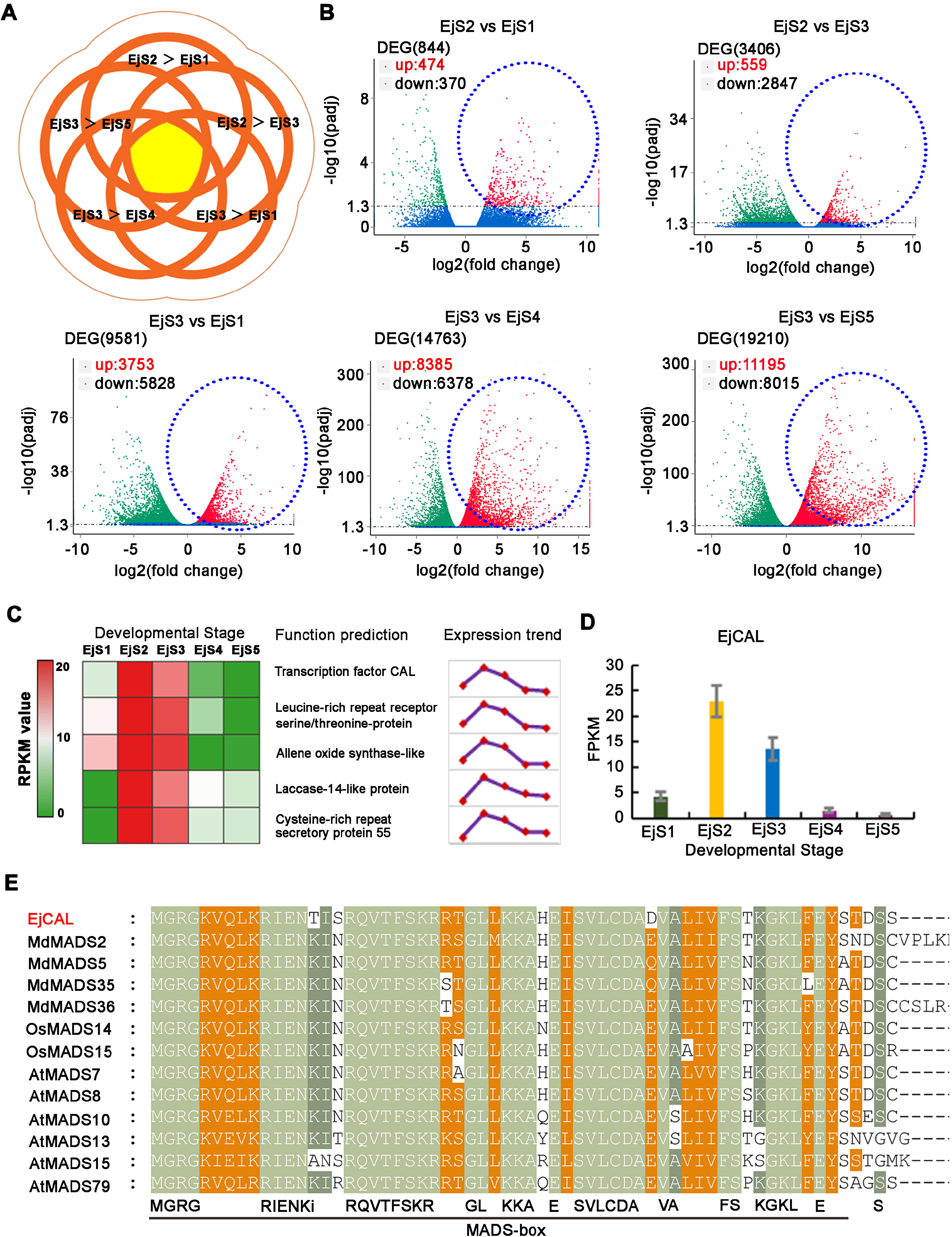
Screening process to identify a transcription factor gene using RNA-seq data from the five stages of flower bud differentiation (FBD). **A,** Gene screening criteria (EjS2 > EjS1, EjS2 > EjS3, EjS3 > EjS1, EjS3 > EjS4, and EjS3 > EjS5) diagram. **B,** Volcano plots of differentially expressed genes based on RNA-seq data. EjS2 > EjS1: 474 genes, EjS2 > EjS3: 559 genes, EjS3 > EjS1: 3753 genes, EjS3 > EjS4: 8385 genes, and EjS3 > EjS5: 11195 genes. **C,** Five candidate genes, comprising one transcription factor (TF), designated *EjCAL*, and four functional genes, were selected. **D,** *EjCAL* expression (highest in the EjS2 stage and about five times higher than in the EjS1 stage). **E,** EjCAL sequence analysis showing its typical MADS-box domain.

**Figure 4.**
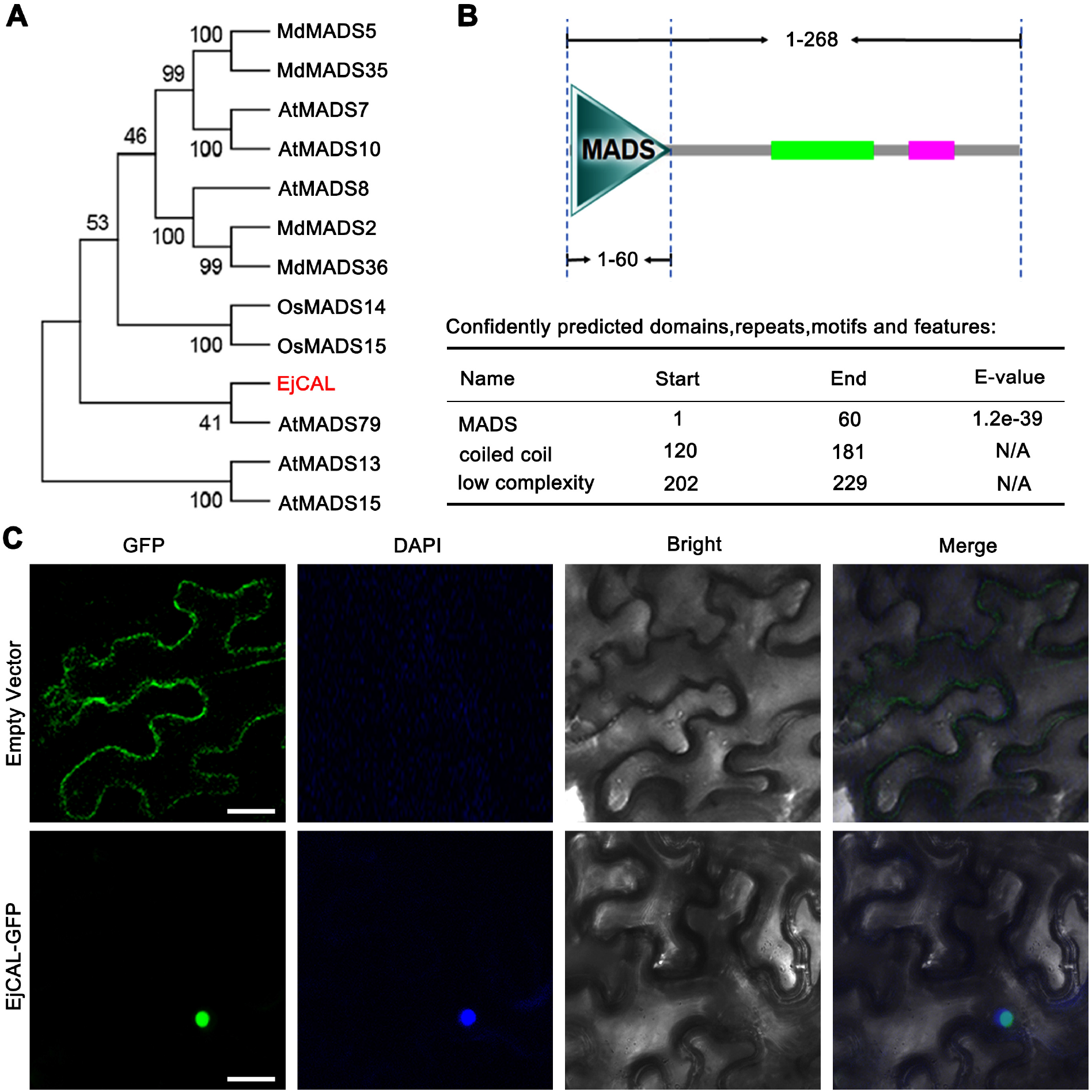
Neighbor-joining tree analysis and subcellular localization of the transcription factor (TF), EjCAL. **A,** Neighbor-joining tree analysis of EjCAL indicating that it is similar to AtMADS79 and belongs to the MADS-box TF family. **B,** It has an N-terminal 60-aa MADS-box domain, a coiled-coil domain, and a low-complexity domain. **C,** Subcellular localization of EjCAL in tobacco epidermal cells. Bars = 20 μm.

### FBD was promoted in *EjCAL*-overexpressing transgenic tobacco

To verify the role of EjCAL in FBD, we constructed a pCAM1300-*EjCAL* vector. Two *EjCAL*-overexpressing (OE) transgenic lines were obtained from 20 initially selected transgenic lines. After selfing, the T2 generation of the two lines (designated EjCAL-OE-#2 and *EjCAL*-OE-#5) had *EjCAL* expression that was 5–8-fold greater than the control lines (empty vector) (Figure 5). We observed the whole process of FBD in the transgenic lines. FBD clearly occurred earlier in EjCAL-OE-#2 and EjCAL-OE-#5 compared to control lines, with flower buds occurring 9 days in advance. The mean number of flower buds and flower bud stage ratio (EjS2–EjS5 stage buds/total buds) were also higher in EjCAL-OE-#2 and EjCAL-OE-#5 compared to control lines. These results indicated that *EjCAL* overexpression promotes early FBD in transgenic tobacco.

**Figure 5.**
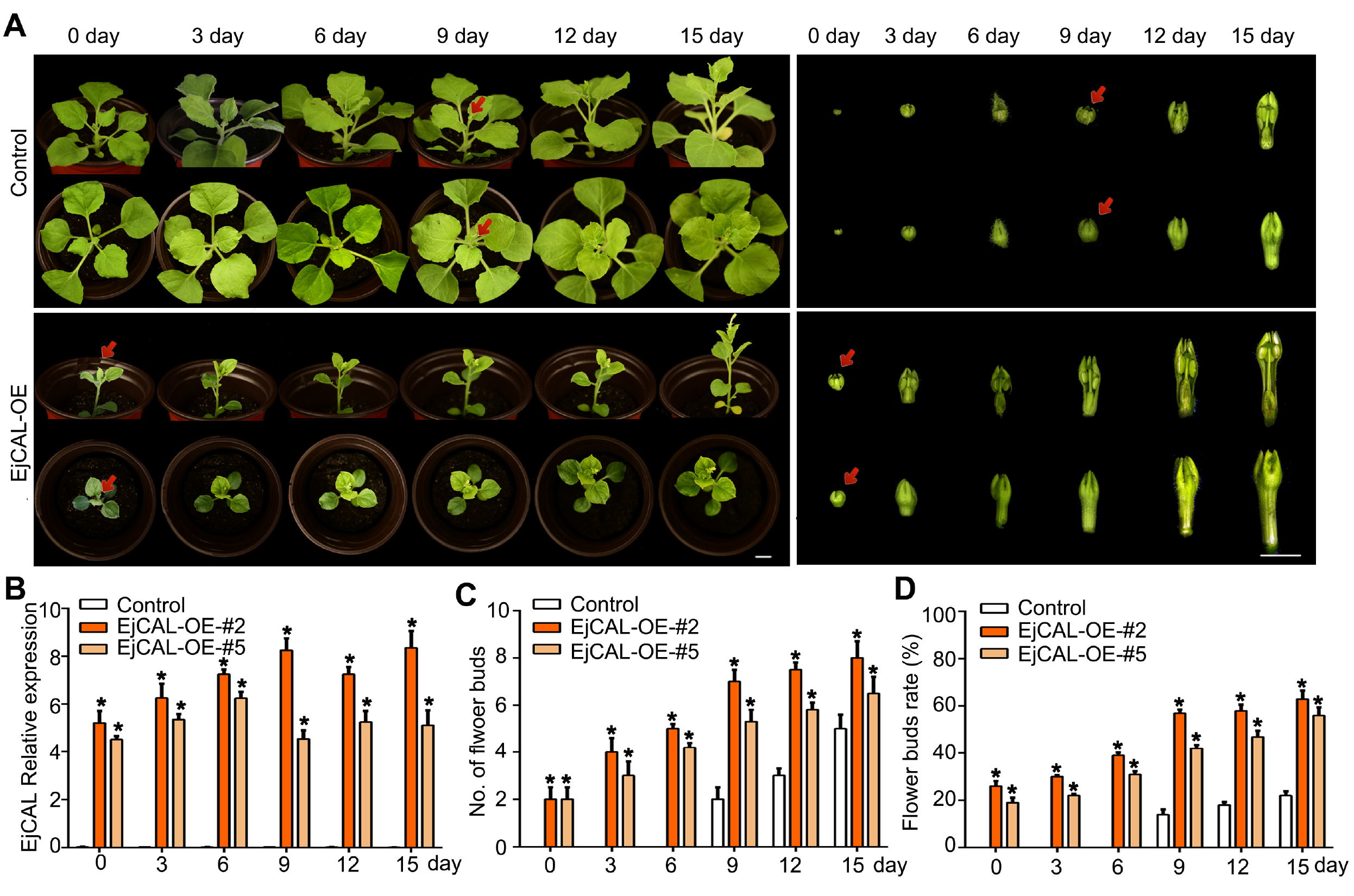
*EjCAL*-overexpressing transgenic tobacco confirmed that EjCAL promotes flower bud differentiation (FBD). **A,** Top-down views and images of individual flower buds of *EjCAL*-overexpressing transgenic tobacco and controls in the critical stages of FBD on days 0, 3, 6, 9, 12, and 15. **B,** Relative *EjCAL* expression in two *EjCAL*-overexpressing transgenic lines (obtained from 20 initially selected transgenic lines; selfing occurred to create the T2 generation of two selected lines designated *EjCAL*-OE-#2 and *EjCAL*-OE-#5). **C,** Number of flower buds in control, *EjCAL*-OE-#2, and *EjCAL*-OE-#5 over 15 days. **D,** Ratio of EjS2-EjS5 stage buds/total buds in control, *EjCAL*-OE-#2, and *EjCAL*-OE-#5 over 15 days. Data are expressed as mean±SEM of n=3 biological replicates. *P<0.05 (Student’s t-test).

### EjERF12 binding to the *EjCAL* promoter is involved in sorbitol-mediated FBD

To further study the mechanism of *EjCAL* in FBD, the *EjCAL* promotor was used as the bait to screen a Y1H library of loquat EjS2 flower buds. Twenty-one candidate genes were found and the function of ten of them were predicted (Table S2). We assessed the open reading frame sequences of the ten genes and conducted pairwise interactions with the *EjCAL* promoter. Only five could bind to it and there was one TF, known as ERF12 (Table S2). In further Y1H assays, the *EjCAL* promoter was divided into four parts, designated P (−2035 to −1 bp), P1 (−755 to −1 bp), P2 (−756 to −1231 bp), and P3 (−1231 to −2035 bp). The Y1H assays showed that only P and P2, which contained the ERF12 binding site (GTCGG), could bind to EjERF12 (Figure 6A). Moreover, we also designed four primer pairs (S1–S4) to perform ChIP-PCR and found that the S1 pair, which corresponded to the P2 region, confirmed the binding relationship between ERF12 and the *EjCAL* promoter (Figure 6B). For S1, the fold enrichment was significantly increased, indicating that the binding site was in P2. Thereafter, the GUS reporter system was used to confirm the function of EjERF12. The result showed that EjERF12 increased GUS activity when it was driven by the *EjCAL* promoter (Figure 6).

**Figure 6.**
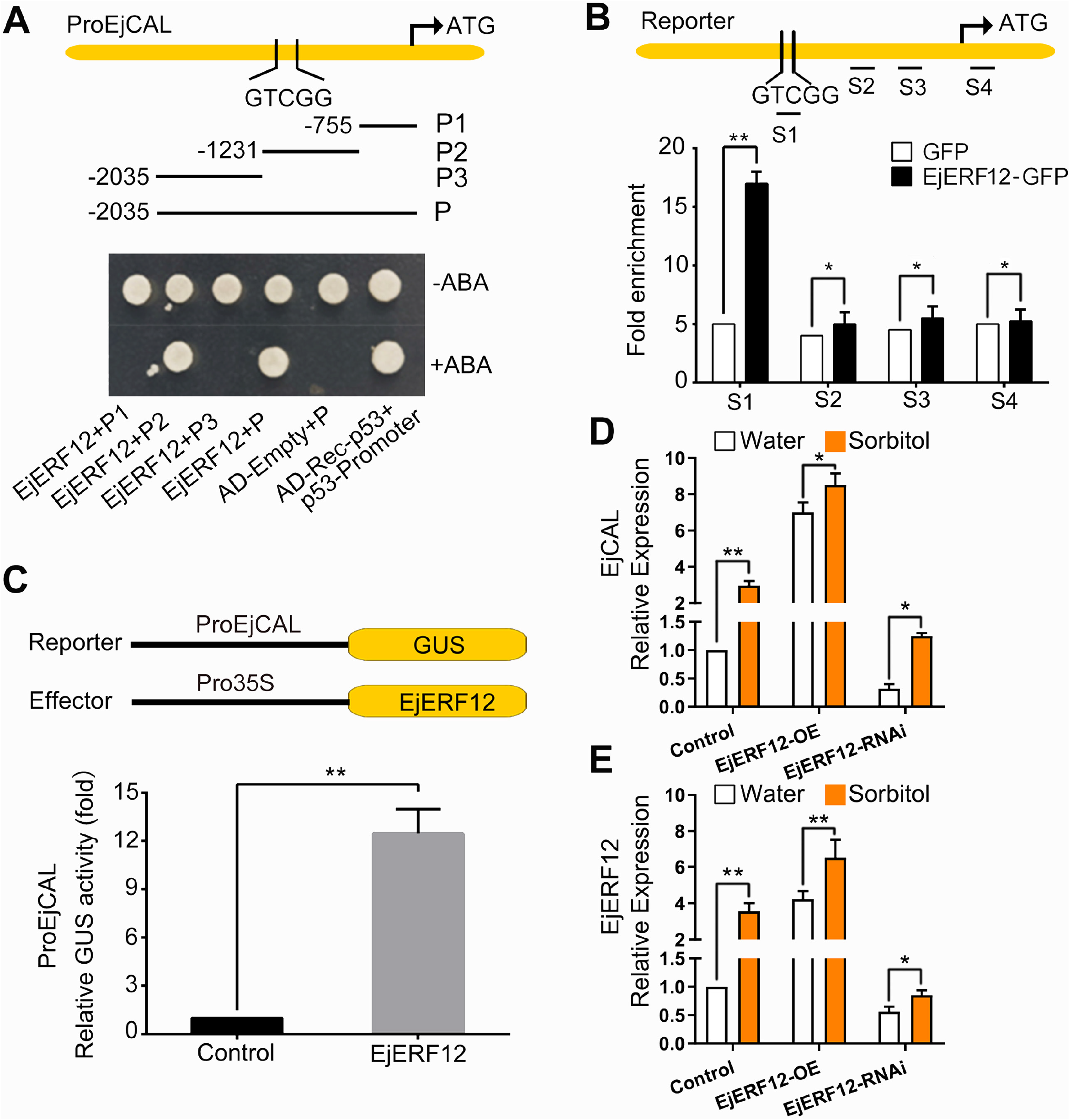
EjERF12 promotes the transcription of *EjCAL*, which is involved in sorbitol-mediated flower bud differentiation (FBD). **A,** Yeast one-hybrid assays showing that EjERF12 binds to the *EjCAL* promoter (*ProEjCAL*). This promoter was divided into four parts (P, P1, P2, and P3). Aureobasidin A (AbA, 200 ng ml^-1^) was used as the screening marker. Rec-p53 + P53-promoter acted as the positive control, as this interaction has been previously confirmed. Empty vector + *ProEjCAL* promoter (P, 2035 bp) acted as the negative control. **B,** ChIP-PCR showing the *in vivo* binding of EjERF12 to the *EjCAL* promoter (using primer pairs S1-S4) in *EjERF12-GFP*-overexpressing and *GFP*-overexpressing (negative control) loquat. Cross-linked chromatin samples were extracted from loquat and precipitated using an anti-GFP antibody, and qPCR was then used to quantify the DNA. The ChIP assay was repeated three times and the enriched DNA fragments in each assay were used as one biological *EjCAL* replicate for qPCR. **C,** GUS activity assay showing that EjERF12 binds to the *EjCAL* promoter. The EjERF12 effector vector and the *ProEjCAL* promoter-containing reporter vector were transfected into wild tobacco (*Nicotiana benthamiana*) leaves. Three independent transfection experiments were performed. **D, E,** *EjCAL* and *EjERF12* expression in control, *EjERF12*-overexpressing, and *EjERF12*-suppression leaves in water and sorbitol spraying groups. *P<0.05, **P<0.01 (Student’s t-test).

After we confirmed the regulation relationship between EjERF12 and *EjCAL*, the involvement of sorbitol was assessed. We overexpressed or suppressed *EjERF12* in the leaves, and then detected *EjERF12* and *EjCAL* expression. Both were upregulated after *EjERF12* overexpression and downregulated after using RNAi against *EjERF12*. After applying sorbitol, both *EjERF12* and *EjCAL* were more highly upregulated compared with those applying water. The results indicated that *EjERF12* and *EjCAL* are involved in sorbitol-mediated regulation in loquat.

### Downstream genes might be involved in sorbitol-mediated FBD

To further study the mechanism of sorbitol-mediated FBD, genes downstream of *EjCAL* were identified, which might regulate parts of FBD. Sorbitol, as an important primary metabolite, may play a signaling role in FBD. When this signal is transmitted *in vivo*, what is the response? Based on RNA-seq data, we found that genes such as *EjUF3GaT1, EjGEF2*, and *EjADF1* may be involved in FBD. Although the expression trends from EjS1 to EjS3 of the three genes met the original screening criteria (EjS2 > EjS1, EjS2 > EjS3, and EjS3 > EjS1), this was not the case for the screening criteria regarding EjS4 to EjS5 (EjS3 > EjS4 and EjS3 > EjS5). After sorbitol treatment, these genes were all significantly upregulated in the EjS2 stage, like for *ERF12* and *EjCAL*. In contrast, the expression of these three genes did not significantly change after either glucose or sucrose treatment, indicating that they might be regulated by sorbitol (Figure 7A). Moreover, the three genes are closely related to hyperoside biosynthesis and signal transduction, e.g., *EjUF3GaT1*, a glycosyltransferase gene, controls hyperoside biosynthesis. Therefore, we explored the changes in hyperoside content during sorbitol-mediated FBD in loquat. Our HPLC results showed that the hyperoside content in loquat flower buds was significantly upregulated after sorbitol treatment. There was little change after glucose or sucrose treatment (Figure 7B).

**Figure 7.**
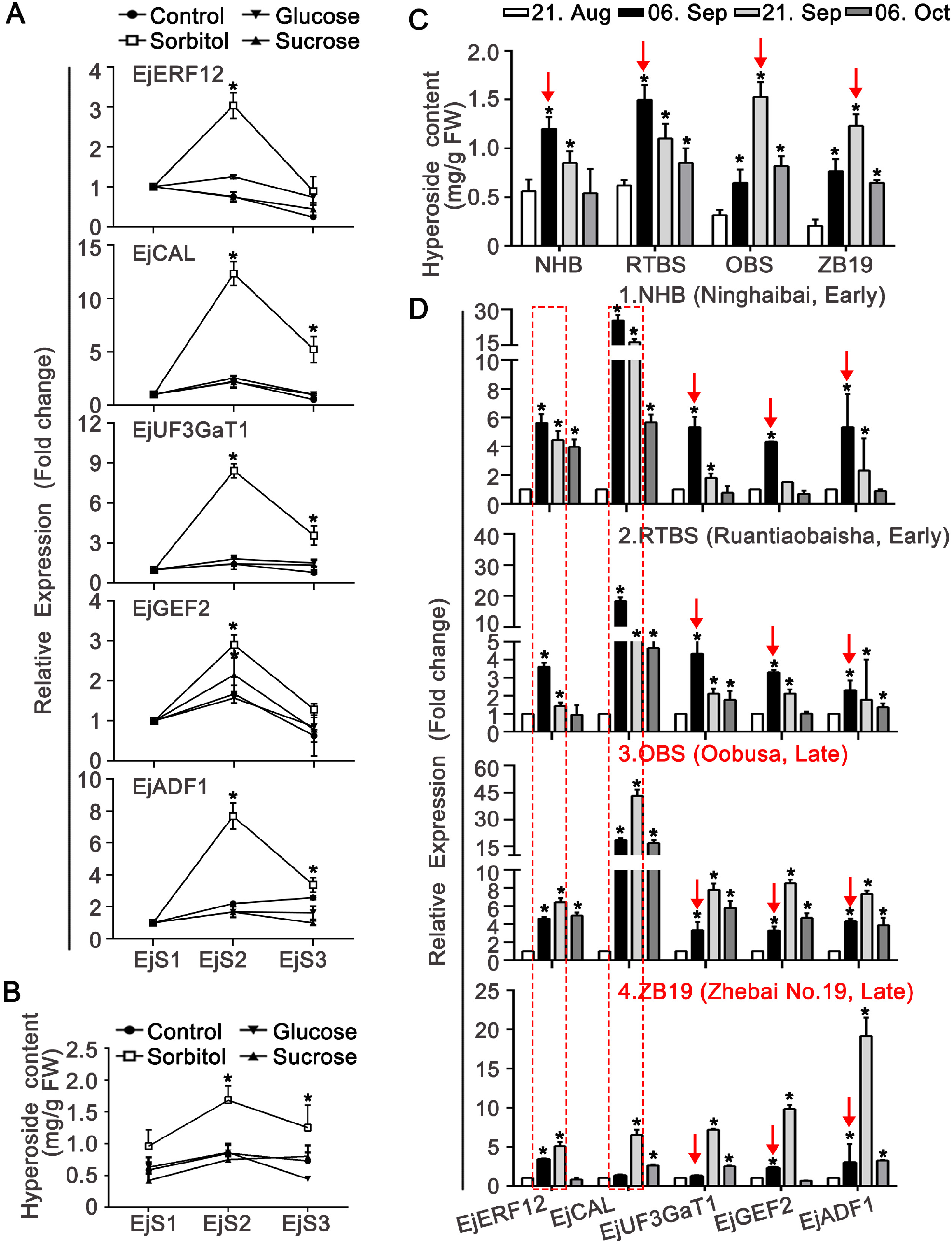
Relative expression of downstream genes involved in flower bud differentiation (FBD), and hyperoside content, after sorbitol, glucose, and sucrose treatments. **A,** Relative expression of *EjERF12, EjCAL*, and key downstream hyperoside biosynthesis pathway genes (*EjUF3GaT1, EjGEF2*, and *EjADF1*) after sorbitol, glucose, and sucrose treatments in the EjS1, EjS2, and EjS3 stages during FBD. **B,** Hyperoside content in the EjS1, EjS2, and EjS3 stages during FBD. **C, D,** Hyperoside content and relative expression of *EjERF12, EjCAL, EjUF3GaT1, EjGEF2*, and *EjADF1* in two early-maturing (NHB and RTBS) and two late-maturing (OBS and ZB19) loquat varieties on August 21, September 6, September 21, and October 6. *P<0.05, **P<0.01 (Student’s t-test).

We then assessed the expression trends of *EjERF12* and *EjCAL* and downstream genes (*EjUF3GaT1, EjGEF2*, and *EjADF1*) and hyperoside content in two early-flowering loquat varieties, NHB (Ninghaibai) and RTBS (Ruantiaobaisha), and two late-flowering loquat varieties, OBS (Oobusa) and ZB19 (Zhebai no. 19) (Figure 7C). *EjERF12* and *EjCAL* expression increased by September 6 and then decreased by September 21 in the two early-flowering varieties, but gradually increased between September 6 to 21 and then decreased by 6 October in the two late-flowering varieties. The expression of the downstream genes (*EjUF3GaT1, EjGEF2*, and *EjADF1*) peaked around September 6 in the early-flowering varieties, and around September 21 in the late-flowering varieties (Figure 7D). Additionally, the hyperoside content peaked early, around September 6, in the early-flowering varieties and later, around September 21, in the late-flowering varieties (Figure 7C).

### Hyperoside biosynthesis was controlled by EjCAL, which might be involved in the sorbitol signaling pathway

Based on the previous results, sorbitol may be a signaling factor that regulates hyperoside biosynthesis, but the molecular mechanism required further verification. First, Y1H assays confirmed that EjCAL can bind to the *EjUF3GaT1* promoter (Figure S1). For the Y1H assays, the *EjUF3GaT1* promoter was divided into four parts, designated P (−1563 to −1 bp), P1 (−456 to −1 bp), P2 (−457 to −1023 bp), and P3 (−1024 to −1563 bp). The Y1H assays showed that only P and P1, which contained the EjCAL binding site (TGAGTTAG; between −1 to −456 of the *EjUF3GaT1* promoter), could bind to the EjCAL protein (Figure S1A). Moreover, we designed four primer pairs (S1-S4) to perform ChIP-PCR, and found that the S1 pair, which corresponded to the P1 region, also confirmed the binding relationship between EjCAL and the *EjUF3GaT1* promoter (Figure S1B). Furthermore, *UF3GaT1* expression was found to be higher in the *EjCAL*-OE-#2 and EjCAL-OE-#5 transgenic tobacco than the control. There were similar results for *GEF2* and *ADF1* (Figure 8B). Lastly, HPLC showed that the hyperoside content was significantly higher in the two transgenic lines than in the control (Figures 8A and 8B).

**Figure 8.**
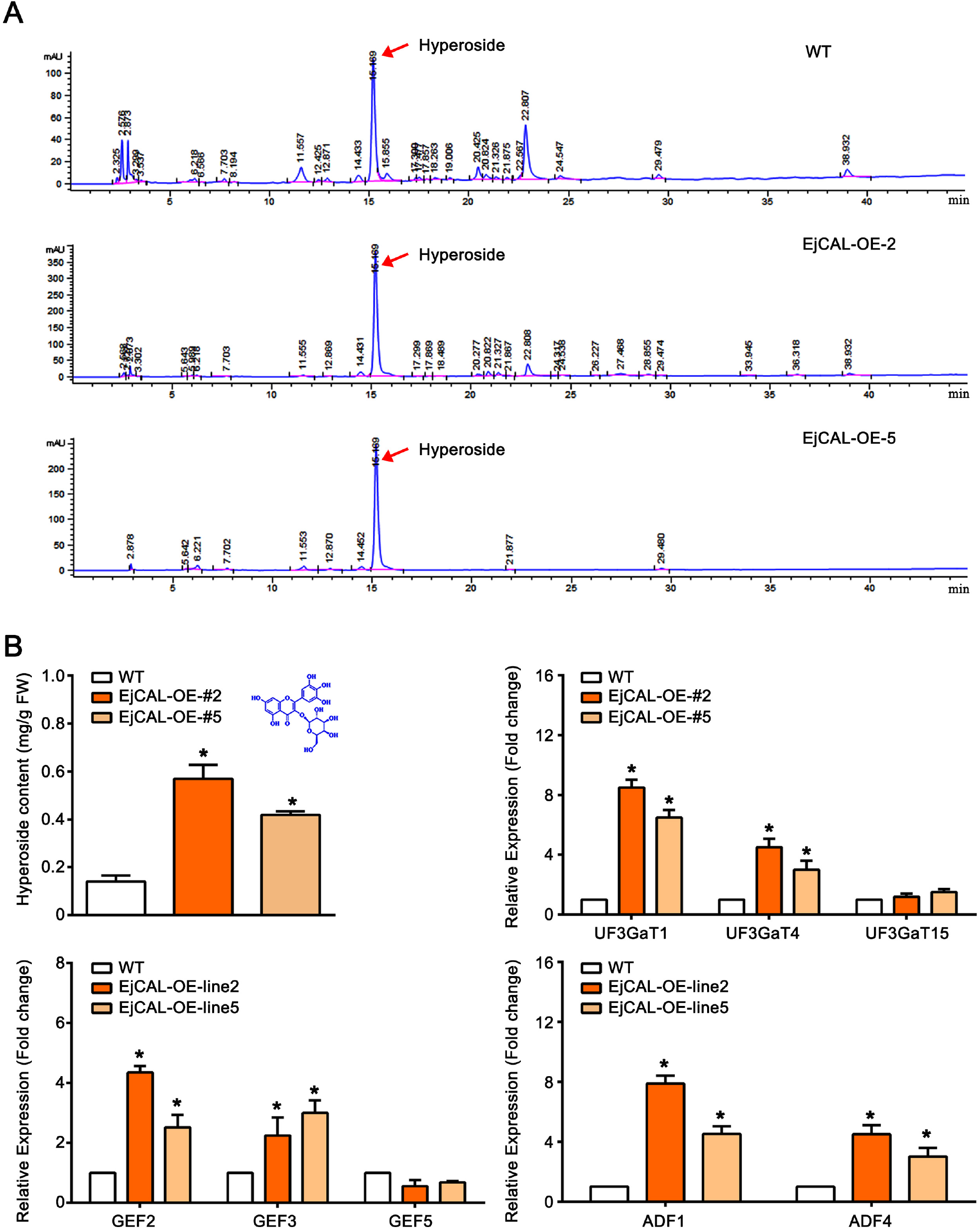
Changes in hyperoside content in transgenic tobacco flower buds. **A,** High-performance liquid chromatography (HPLC) outputs for wildtype, EjCAL-OE#2, and EjCAL-OE#5 transgenic tobacco flower buds. The peak at 16.169 min represents hyperoside. **B**, Hyperoside content and relative expression of the main hyperoside biosynthesis pathway genes (*EjUF3GaT1, EjUF3GaT4*, and *EjUF3GaT15*) and regulatory genes downstream of hyperoside biosynthesis (*EjGEF2, EjGEF3, EjGEF5, EjADF1*, and *EjADF4*) in wildtype, EjCAL-OE#2, and EjCAL-OE#5 transgenic tobacco flower buds. Data are expressed as mean±SEM of n=3 biological replicates. Student’s t-test: *P<0.05 (Student’s t-test).

**Figure 9.**
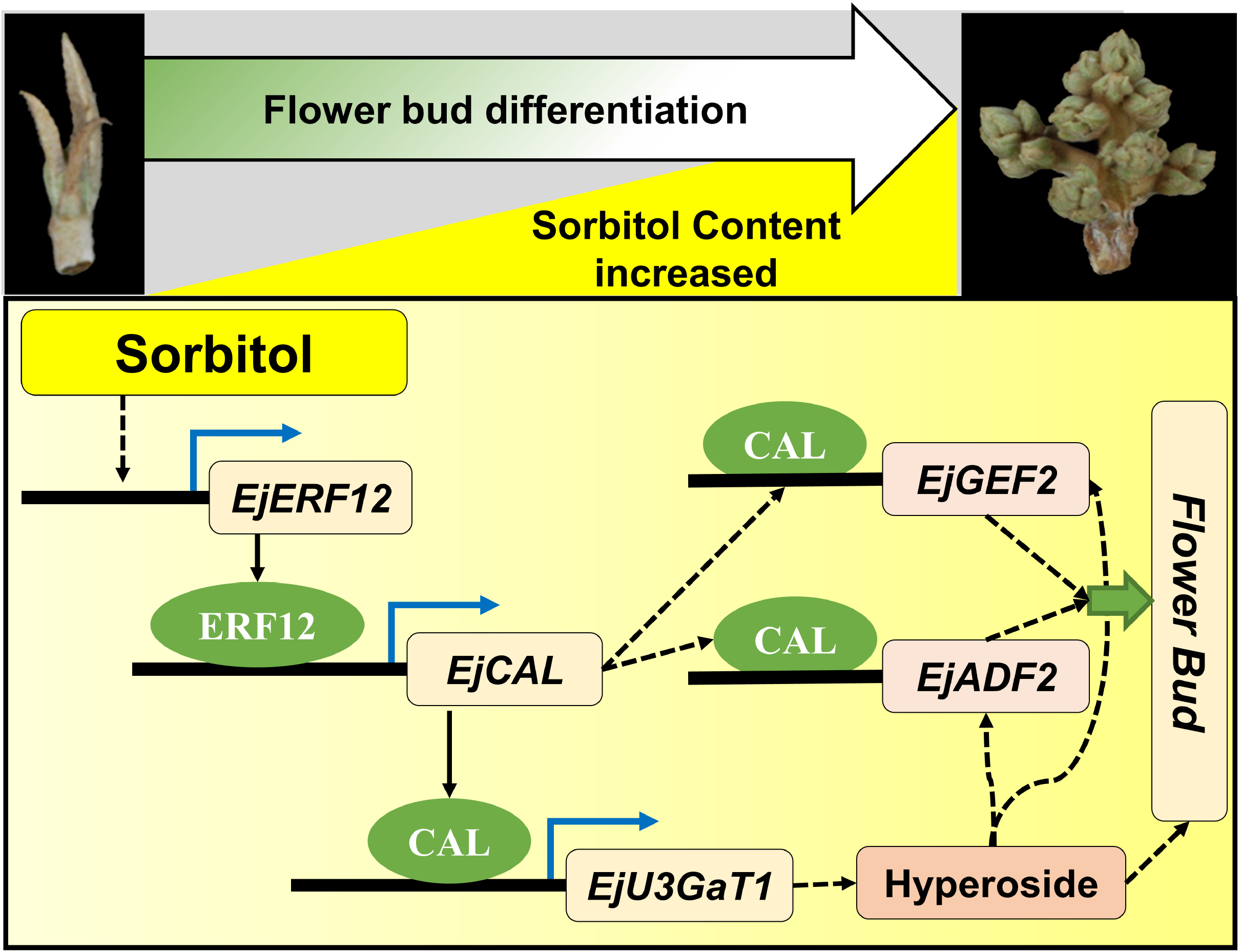
Proposed model of the role of sorbitol accumulation in early flower bud differentiation (FBD) and increased flower bud stage ratio. Sorbitol induces the transcription factor EjERF12 to upregulate *EjCAL* by binding to the GTCGG site in the *EjCAL* promoter. The upregulated EjCAL TF binds to the promotor of *EjUF3GaT1*, a glycosyltransferase that is the final limiting enzyme in hyperoside biosynthesis. The resultant accumulated hyperoside upregulates the downstream reproductive development related genes *EjGEF2* and *EjADF*1. On the other side, sorbitol could also induce the accumulation of hyperoside, and then induce FBD in loquat. The results indicates that sorbitol does play an improtant role in the development of loquat.

## Discussion

### Sorbitol is essential for FBD in loquat

As an important photosynthetic product, sorbitol is widely found in plants, especially in Rosaceae fruit trees. However, there have been few reports on the effects of sorbitol signals on plant development; most related studies have focused more on other sugar signals (e.g., the role of fructose in regulating flower development in *Vernonia herbacea* (Vell.) Rusby (Portes and Carvalho 2006) and the role T6P in regulating FBD in *A. thaliana* (Carvalho and Dietrich, 1993; Wahl et al., 2013). Many studies on fruit trees have found that sorbitol accumulation and changes in FBD or fruit development are closely related, suggesting that sorbitol plays an important regulatory role (Ito et al., 2002). During flower development and pollen tube growth in apple, sorbitol regulates a key TF, MYB39L, which in turn controls downstream developmental genes (Meng et al., 2018). Recent research has also shown that sorbitol controls the pollen tubule transporter HT1.7 by regulating the transport of other sugars, such as sucrose and fructose, into the pollen tube, thereby controlling its growth (Li et al., 2020). These studies suggest that sorbitol plays an important signaling role in floral organ development. These findings are similar to our findings on the regulatory role of sorbitol in FBD in loquat. We found that sorbitol accumulated during the critical period of FBD. Based on the FBD phenotypic data and the sorbitol spraying experiment, we speculated that sorbitol plays an important role in the occurrence and development of FBD.

### EjCAL plays a key role in sorbitol-mediated FBD

We screened transcriptome data from different stages of FBD and obtained a TF, designated *EjCAL*, which belongs to the MADS-box family (Figure 3). As members of the MADS-box family, *CAL* genes encode MIKC-type MADS-box domain-containing TFs and participate in plant development (Kempin et al., 1995). In *Arabidopsis, AP1* and *CAL* are a pair of duplicate genes that have redundant function regarding controlling the initiation of flower development (Bowman et al., 1993; Goslin et al., 2017). Interestingly, *AP1* and *CAL* have diverged considerably in terms of temporal, spatial, and level of expression by gaining and losing cis-regulatory elements (Ye et al., 2016). However, unlike *AP1*, whose function has been widely reported, information on *CAL* is very limited, and there have been no studies on *EjCAL*. We found that *EjCAL* expression was highly correlated with FBD phenotypic data, indicating its critical role in FBD initiation. Using *EjCAL*-overexpressing transgenic tobacco plants, we confirmed that *EjCAL* can initiate early flower bud development (Figure 5). It has not been previously reported that sorbitol plays a signaling role in regulating *CAL* and affecting FBD. The important role of sorbitol in fruit tree development and fruit quality, and particularly FBD, is an important research direction. During flower development, the changes in *EjCAL* expression were consistent with sorbitol accumulation (Figure 3). It was also found that *EjCAL* expression changed significantly after spraying sorbitol (Figure7). Our results have given us a clearer understanding of FBD in loquat. Sorbitol, as a sugar signaling substance, may be used as a marker of FBD regulation, and *EjCAL* is likely to be a key hub gene for regulating FBD.

### ERF12 and many downstream genes might be involved in sorbitol-mediated FBD

The ethylene response factor (ERF) family is a family of plant-specific TFs. In addition to participating in the response to and regulation of ethylene signaling (Zhang et al., 2009; Li et al., 2016), ERF family members also play a variety of roles in flower bud transformation (Zhu et al., 2021). Ethylene is an important factor affecting female FBD and ovary development in plants (García et al., 2020). The TF ERF12 promotes early FBD and is an important regulatory factor in bud breaking in transgenic poplar (Yoshikawa et al., 2014). However, there are few studies on the function of *ERF* in sugar signaling.

To clarify the upstream and downstream regulatory network related to the core sorbitol-regulated TF *EjCAL*, we carried out Y1H screening involving the *EjCAL* promoter and identified EjERF12. EjERF12 may be a key part of the sorbitol signaling cascade that leads to *EjCAL* upregulation and, in turn, regulation of downstream genes by *EjCAL*, or it may act as a cross point between sorbitol signaling and other hormone signals.

We also verified the transcriptome data, analyzed the related functional genes, and selected and verified the functional genes that may be regulated by *EjCAL* and may play a role in FBD, including *EjUF3GaT1, EjGEF2*, and *EjADF1. UF3GaT* is a key synthase gene in flavonoid metabolism, playing a key role in the biosynthesis and accumulation of flavonoids such as hyperoside (Yang et al., 2020a; 2020b). Rho-type guanine nucleotide exchange factor (*RhoGEF*) encodes a key factor that regulates the activity of Rho GTPase, which is involved in processes such as cytoskeleton regulation, transcription, and membrane vesicle transport in eukaryotic cells, acting as a molecular switch. It is a gene that plays a vital role in sensory signal regulation, signal transduction, regulation of cell growth and development, and immune response (Kashyap et al., 2019). Lastly, *ADF* encodes a microfilament-associated protein that is closely related to cell growth and development (Zheng et al., 2013; Tian et al., 2009). These three downstream functional genes may play different roles in FBD, and all appear to play important roles in FBD in loquat.

### Hyperoside may be regulated by sorbitol signaling and act on downstream genes

In addition to the key genes in FBD being affected by the sorbitol-*EjERF-EjCAL* pathway, we also noted significant changes in *EjUF3GaT1*, suggesting that sorbitol signaling may regulate the accumulation of the flavonoid hyperoside. Regarding this hypothesis, we found four relevant pieces of evidence: (1) sorbitol accumulation in EjS1–EjS3 during FBD in loquat was accompanied by the accumulation of hyperoside; (2) *EjUF3GaT1* was upregulated and hyperoside accumulated after sorbitol spraying; (3) the hyperoside content in *EjCAL*-overexpressing transgenic tobacco was significantly higher than that in the control; and (4) the TF EjCAL could bind to the promoter region of *EjUF3GaT1* and regulate its expression. This evidence suggested that sorbitol may act as an upstream signal to regulate hyperoside accumulation via EjCAL. In recent years, there has been increasing evidence that flavonoids not only significantly accumulate during plant reproductive development, but also exhibit important functions during development and stress resistance (Yang et al., 2020b; Li et al., 2021; Górniak et al., 2018). In okra, hyperoside regulates the flowering time, pollen tube growth, and seed set rate (Yang et al., 2020b), suggesting that hyperoside may have an important signaling function. In this study, we identified an important role of sorbitol in hyperoside accumulation, which may involve a synergistic effect between sorbitol signaling and flavonoid metabolism or flavonoid signaling, and the related findings provide important evidence for further in-depth study of the roles of sorbitol and flavonoid signaling in FBD.

## Conclusions

Our research indicated that sorbitol significantly accumulated during FBD from the EjS1 stage to the EjS2 stage, and sorbitol then significantly decreased by the time of FBD completion in the EjS3 stage. Transcriptome data showed that a MADS-box TF designated *EjCAL* was highly correlated with the FBD phenotypic data. Y1H and ChIP-PCR assays confirmed that EjERF12 could bind to the *EjCAL* promoter and regulate its expression. We also found that sorbitol-mediated FBD involved the accumulation of an important developmental flavonoid, hyperoside, as EjCAL binds to the *EjUF3GaT1* promoter, a key gene for hyperoside biosynthesis, and thereby promotes hyperoside accumulation. Additionally, in the EjS2 stage compared to the EjS1 stage, it was found that *EjGEF2* and *EjADF1* (reproductive development-related genes downstream of hyperoside) were obviously upregulated. Two early- and late-flowering varieties of loquat showed that *EjERF12* and *EjCAL* expression was closely related to flower development stage. In summary, the molecular mechanism regarding how sorbitol regulates FBD through *EjERF12* and *EjCAL* was confirmed.

## Materials and methods

### Plant materials

The experiment was conducted in the Base Orchard of the Zhejiang Academy of Agricultural Sciences (Jiaxing, Zhejiang Province, China) in 2018–2020. Loquat (*Eriobotrya japonica* Lindl.) cv. ‘Ninghaibai’ (NHB), ‘Ruantiaobaisha’ (RTBS), ‘Oobusa’ (OBS), and ‘Zhebai no. 19’ (ZB19) were used in this study. The trees were 10 years old, were maintained using standard horticultural practices, and were treated with standard disease and insect control measures.

### Methods

#### Flower bud morphological observation

The following five stages of flower bud development in loquat were investigated: physiological FBD (EjS1), morphological FBD (EjS2), panicle development (EjS3), florescence development (EjS4), and flowering (EjS5) stages. Each flower bud was cut into 0.5-cm slices at the best position and then immediately placed into a prepared fixative solution (5 mL formalin, 5 mL glacial acetic acid, 90 mL 70% ethanol, and 5 mL glycerol). The fixed tissues were dehydrated, cleared, embedded in paraffin, modified section of wax block, adhesive sheet spreading, dyed, sealed, and then placed under a biological microscope for morphological observation.

#### Determination of main metabolic factors

The above five stages of the flower buds of the white-flesh loquat variety NHB were used to analyze the main metabolites (soluble sugar, starch, glucose, fructose, sucrose, and sorbitol).

Soluble sugar contents were determined as previously described (Song et al., 2020). Briefly, 0.1g of flower buds was placed in a boiling water bath for 1 h. After the extract was naturally cooled, 0.2 mL of the extract was taken, and 5 mL concentrated sulfuric acid, 1.8 mL distilled water and 0.5 mL anthrone were added. The mixture was shaken for 5 min and heated at 100°C for 1 min, and the absorbance at 620 nm was then assessed. D-glucose was used to create a standard curve.

To assess the starch content, 0.1 g of flower buds at different stages was ground up, followed by adding 2 mL anhydrous ethanol and 10 mL 0.5 mol/L NaOH and then heating, dissolution, and cooling. Thereafter, 5 mL of the above solution was diluted with distilled water and adjusted to about pH 3. Next, 1 mL iodine reagent was added followed by adding distilled water to make up the volume to 100 mL. The absorbance at 620 nm was then measured (with 1 mL iodine solution in 99 mL distilled water used as the blank control), and the starch content was calculated using a standard curve.

Agilent 7890B gas chromatography coupled with 7000D mass spectrometer (Agilent Technology, Palo Alto, CA, USA) was used to simultaneously assess the glucose, fructose, sucrose, and sorbitol contents, as previously described (Meng et al., 2018) with some modifications. Briefly, each frozen flower bud sample was crushed, sugars were extracted from 100mg of power in the solution of methanol: isopropanol: water (3:3:2 V/V/V). The extracts was centrifuged and the supernatants were mixed with ribitol added as an internal standard, and dried under vacuum without heat and the residue was derivatized sequentially with methoxyamine hydrochloride and BSTFA. The samples and prepared standards for each sugar were then separated and analyzed by GC-MS/MS.

#### Sorbitol, sucrose, and glucose spraying

The effects of sorbitol, sucrose, and glucose on the NHB loquat variety were studied using a completely randomized design. Nine single-tree replicates per treatment were sprayed with 1.0% sorbitol, 2.0% sucrose, 2% glucose, or water (control) (until runoff occurred) at the physiological FBD stage (EjS1) on July 17, respectively. After spraying for 40, or 80 d, the flower bud stage ratios (EjS2–EjS5 stage buds/total buds) were assessed. In addition, the flower buds in the EjS1, EjS2, and EjS3 stages were harvested for further analysis.

#### Quantitative real-time (qRT)-PCR of flower buds

Flower buds at various developmental stages (EjS1, EjS2, and EjS3) as well as the flower buds of two early-flowering varieties (NHB and RTBS) and two late-flowering varieties (OBS and ZB19) harvested on August 21, September 6, September 21, and October 6 in 2020, underwent qRT-PCR. The total RNA of each flower bud was extracted using the cetrimonium bromide (CTAB) method. cDNA was synthesized using PrimeScript RT Reagent with a FastQuant RT SuperMix kit (Tiangen Biotech Company, Beijing, China). The cDNA quality of three biological replicates was assessed using a NanoDrop spectrophotometer (ThermoScientific Inc., Waltham, MA, USA). qRT-PCR analysis of the cDNAs was then conducted using Fast Fire qPCR PreMix (SYBR Green, Tiangen Biotech Company, Beijing, China) on a CFX Connect Real-Time PCR Detection System (Bio-Rad, CA, USA). The primers of the genes analyzed by qRT-PCR are listed in Table S1. The expression data were analyzed using the 2^-ΔΔCt^ method.

#### Transcriptome sequencing of flower buds

The total RNA was extracted from flower buds at various developmental stages (EjS1, EjS2, EjS3, EjS4, and EjS5) via the CTAB method and used to generate cDNA libraries. An Illumina NovaSeq 6000 platform (Illumina Inc., San Diego, CA, USA) was used for transcriptome sequencing. To improve the assembly accuracy, we removed ends with <q20 quality from the reads. Reads with <50 bp, >0.2 error rate, or >10% ambiguous (N) bases were also excluded. The remaining high-quality reads were consistent with the most recent loquat genome. We only retained the uniquely mapped reads. Differentially expressed genes (DEGs) in flower buds at different developmental stages (EjS2 vs. EjS1, EjS2 vs. EjS3, EjS3 vs. EjS1, EjS3 vs. EjS4, and EjS3 vs. EjS5) were identified using the edgeR package. We built a model to reduce the explanatory factors into a single column for the intercept, estimated the dispersion from the model using the GLMCommonDisp function with the parameters method, and then assessed the DEGs using the exact test with the estimated dispersion.

#### Sequence alignment and annotation of EjCAL in loquat

The EjCAL protein sequence was used as a query sequence, and 4 MADS-box protein sequences in apple (from the apple genome sequences in the Genome Database of Rosaceae [GDR]; https://www.rosaceae.org/), 6 in *Arabidopsis* (from The Arabidopsis Information Resource [TAIR]; http://www.Arabidopsis.org/, accessed on October 10, 2016), and 2 in rice (from the National Center for Biotechnology Information [NCBI] website) were returned. Structural motif annotation was performed using DNAMAN7 software. Phylogenetic analysis was conducted using MEGA v6.0 (http://www.megasoftware.net/, accessed October 10, 2016) with the neighbor-joining method and bootstrap values calculated using 1000 iterations. The EjCAL sequences were used to query the Simple Modular Architecture Research Tool (SMART) database (http://smart.embl-heidelberg.de/, accessed on October 10, 2016) to determine whether they were predicted to belong to the CAL family.

#### EjCAL subcellular localization analysis

*EjCAL-enhanced green fluorescent protein (eGFP)* was inserted into a pEZS-NL vector. *EjCAL* and *eGFP* were fused and inserted into a pCAMBIA1300 vector under the control of the CaMV35S promoter. The vector was transformed into *Agrobacterium tumefaciens* GV3101, which was centrifuged to obtain the pellet and resuspend it in buffer. Wild tobacco (*Nicotiana benthamiana*) leaves were immersed in the solution (using a 5 mL syringe), incubated in a greenhouse at 22–24°C for 72 h, and then observed under a fluorescence microscope. eGFP was excited using 488 nm light and the nuclei stain 4’,6-diamidino-2-phenylindole (DAPI), which is often used to detect TFs, was excited using ultraviolet light. The detection wavelengths were 500–530 nm for eGFP and 430–460 nm for DAPI.

#### *EjERF12*-overexpressing and *EjERF12*-suppression vector construction

*EjERF12* cDNA was cloned into a pROK2 vector by homologous recombination following Gateway Cloning Protocols using BP Clonase II enzyme mix (Thermo Fisher Scientific). Additionally, an RNA interference (RNAi) vector was constructed using a pFGC5941 vector (BioVector Co., LTD Beijing, China) following the manufacturer’s instructions. The *EjERF12* overexpression and RNAi vectors were separately transformed into *A. tumefaciens* GV3101 by the electric shock method, respectively.

#### Generation of *EjCAL*-overexpressing transgenic tobacco

Transgenic tobacco was obtained according to a previously described method. Briefly, the full-length coding sequence of *EjCAL* was inserted into a pBI121 vector under the control of the CaMV35S promoter and then transformed into tobacco leaf discs using *A. tumefaciens* GV3101. An initial 20 transgenic tobacco lines were selected using 50 mg L^-1^ kanamycin and further confirmed by PCR. Subsequently, the T1 generation seeds were subjected to segregation analysis. Transgenic lines with a chi-square value <1.5 (p ≤ 0.05) were considered to be true transgenic lines. Independently regenerated transgenic lines exhibiting 3:1 segregation ratios of kanamycin resistance to kanamycin sensitivity were selected and the respective T2 generation transgenic plants were propagated in a greenhouse. The primers used are listed in Table S1.

#### Yeast one-hybrid (Y1H) assays

For the Y1H assays, a Matchmaker Gold Y1H System (Clontech) was used following the manufacturer’s instructions. In the first set of Y1H assays, the *EjCAL* promotor was used as the bait to screen a cDNA prey library based on loquat EjS2 flower buds. The *EjCAL* promotor was ligated to a pAbAi bait vector after linearizing the vector using BstBI and then transformed into the Gold yeast strain and selected on an SD/-Ura plate.

In the second set of Y1H assays, *EjCAL* promoter fragments (P [−2035 to −1 bp], P1 [−755 to −1 bp], P2 [−756 to −1231 bp], and P3 [−1231 to −2035 bp]) or *EjUF3GaT1* promoter fragments (P [−1563 to −1 bp], P1 [−456 to −1 bp], P2 [−457 to −1023 bp], and P3 [−1024 to −1563 bp]) were each ligated to a pAbAi bait vector after linearizing the vector using BstBI. These vectors were then separately transformed into the Gold yeast strain and selected on an SD/-Ura plate. Prey vectors were generated by separately inserting the TFs *EjERF12* and *EjCAL* in-frame with the *GAL4* activation domain in a pGADT7 vector.

In the Y1H assays, the prey vectors (with the cDNA prey library, *EjERF12*, or *EjCAL*) were separately transformed (using LiCl-polyethylene glycol) into the Gold yeast strain harboring one of the above mentioned bait vectors. The cDNA prey library transformants and *EjERF12/EjCAL* transformants were grown on SD/-Leu/Aureobasidin A (AbA) or SD/-Ura/AbA plates (200 ng ml^-1^ AbA), respectively, at 30°C for 3–5 days. Vectors from positive clones were isolated from the yeast and then individually transferred into *Escherichia coli* DH5α cells for amplification and sequencing.

#### ChIP-PCR

ChIP-PCR was used to show that EjERF12 binds to the *EjCAL* promoter and that EjCAL binds to the *EjUF3GaT1* promoter (with each experiment using primer pairs S1-S4, which correspond to different parts of the *EjCAL* and *EjUF3GaT1* promoters, as shown in Figures 6B and S2B, respectively) in *EjERF12*- or *EjCAL-GFP*-overexpressing and *GFP*-overexpressing (negative control) loquat.

The *EjCAL* promoter was divided into four parts, designated, as mentioned previously, P (−2035 to −1 bp), P1 (−755 to −1 bp), P2 (−756 to −1231 bp), and P3 (−1231 to −2035 bp). The bioinformatics-predicted (http://bioinformatics.psb.ugent.be/webtools/plantcare/html/#opennewwindow) EjERF12 binding site (GTCGG) was contained in P and P2. The *EjUF3GaT1* promoter was also divided into four parts, designated, as mentioned previously, P (−1563 to −1 bp), P1 (−456 to −1 bp), P2 (−457 to −1023 bp), and P3 (−1024 to −1563 bp). The bioinformatics-predicted EjCAL binding site (TGAGTTAG) was contained in P and P1.

Four pRI101-*GFP-EjERF12* constructs and four pRI101-*GFP-EjCAL* constructs, each with one of the above four regions of the *EjCAL* or *EjUF3GaT1* promoters, were separately transformed into loquat leaves. ChIP assays were then performed using anti-GFP antibody (Transgen Biotech Co., LTD. Beijing, China), with the immunoprecipitated chromatin quantified by qPCR. The primers used are listed in Table S1. Each ChIP assay was repeated three times and the enriched DNA fragments (1 μL) in each ChIP assay were used as one biological replicate for qPCR.

#### β-glucuronidase (GUS) activity assays

A reporter vector containing the *EjCAL* promoter (2035 bp upstream of the start ATG) was prepared. The reporter and EjERF12 effector vectors were transfected into tobacco leaves, and 3 days later proteins were extracted from the leaves and subjected to GUS activity assays. A VersaFluor fluorometer (Bio-Rad) was used to assess the fluorescence of the proteins. GUS activity was determined based on three biological replicates. The primers used are listed in Table S1.

#### Hyperoside content analysis

The hyperoside content in loquat flower buds at different developmental stages and *EjCAL*-overexpressing transgenic tobacco was analyzed. First, 1g of flower buds were dried in an oven at 40–45°C, placed in 100 ml 70% ethanol, homogenized at 15 000 rpm using a high-speed disperser (IKAT18), and centrifuged at 25 000 g. Next, 1 ml of the supernatant was mixed with 0.3 ml 5% sodium nitrite solution for 6 min. Thereafter, 0.3 ml 10% aluminum nitrate was added for 6 min at room temperature, and then 4 ml 4% sodium hydroxide solution was added for 15 min at room temperature. Subsequently, the total volume was brought to 10 ml with 68% ethanol. The absorbance was measured at 510 nm, with 68% ethanol used as the blank control, using a 1260 Liquid Chromatography System (Agilent, Santa Clara, CA, USA) equipped with a Luna C18 column (250×4.6 mm, 5 μm; Phenomenex). The gradient elution conditions, involving 0.1% formic acid as mobile phase A and acetonitrile as mobile phase B, were as follows: 0–45 min, 15–25% B; 45–60 min, 25–35% B. The total run time was 40 min and the flow rate was constant at 1 mL min^-1^. The injection volume of each sample was 20 μL, and the column temperature was maintained at 35°C. Standard was used to quantify the hyperoside content.

#### Statistical analysis

The statistical analysis was performed using SPSS 20.0 (SPSS, Chicago, USA). Using at least three biological replicates, error bars and significant differences between means were determined using Student’s *t*-test.

## Acknowledgements

We would like to thank Dr Yong Yang for skillful technical assistance in generation of transgenic tobacco. We thank Dr Yanna Shi for helpful discussion and critical reading of the manuscript.

## Supplemental data

**Supplemental Table S1.** Primers used for qRT-PCR analysis, Y1H assays, transient ransfection assays, transgenic assays, ChIP-PCR assays, and GUS transactivation assays in this study.

**Supplemental Table S2.** Positive clones obtained in yeast one-hybrid assays using the *EjCAL* promoter as bait.

**Supplemental Figure S1.** EjCAL promotes the transcription of *EjUF3GaT1*, which is involved in hyperoside biosynthesis, during flower bud differentiation (FBD).

